# Cell growth model with stochastic gene expression helps understand the growth advantage of metabolic exchange and auxotrophy

**DOI:** 10.1101/2020.08.26.268771

**Authors:** Dibyendu Dutta, Supreet Saini

**Author notes:** **Corresponding Authors:** Dibyendu Dutta.

## Abstract

During cooperative growth, microbes often experience higher fitness, due to sharing of resources by metabolic exchange and herd protection through biofilm structures. However, the trajectory of evolution of competitive species towards cooperation is not known. Moreover, existing models (based on optimisation of steady-state resources or fluxes) are often unable to explain the growth advantage for the cooperating species, even for simple reciprocally cross-feeding auxotrophic pairs. We present an abstracted model of cell growth that considers the stochastic burst-like gene expression of biosynthetic pathways of limiting biomass precursor metabolites, and directly connects their cellular levels to growth and division using a “metabolic sizer/adder” rule. Our model recapitulates Monod’s law and yields the experimentally observed right-skewed long-tailed distribution of cell doubling times. The model further predicts the growth effect of secretion and uptake of metabolites, by linking it to changes in the internal metabolite levels. The model also explains why auxotrophs may grow faster when provided the metabolite they cannot produce, and why a pair of reciprocally cross-feeding auxotrophs can grow faster than prototrophs. Overall, our framework allows us to predict the growth effect of metabolic interactions in microbial communities and also sets the stage to study the evolution of these interactions.

**Importance:** Cooperative behaviours are highly prevalent in the wild, but we do not understand how it evolves. Metabolic flux models can demonstrate the viability of metabolic exchange as cooperative interactions, but steady-state growth models cannot explain why cooperators grow faster. We present a stochastic model that connects growth to the cell’s internal metabolite levels and quantifies the growth effect of metabolite exchange and auxotrophy. We show that a reduction in gene expression noise explains why cells that import metabolites or become auxotrophs can grow faster, and also why reciprocal cross-feeding of metabolites between complementary auxotrophs allow them to grow faster. Our framework can simulate the growth of interacting cells, which will enable us to understand the possible trajectories of the evolution of cooperation *in silico*.

## Introduction

Unlike the pure cultures in a laboratory, in the wild, microbes live alongside many different species. To survive, they may either compete for limited resources or cooperate. Defying general evolutionary expectations, cooperative communities pervade different ecological niches (1–5), highlighting the purported selective advantage of cooperation (6). Cooperative interactions often involve secretions, ranging from extracellular enzymes, scavenging molecules (like siderophores) (7), to leaked metabolites – either toxic by-products (8, 9) or essential metabolites like amino acids (10–12). Cooperation may also involve living together in a biofilm, which protects the group from the effect of toxins, antibiotics, and predators (13).

To fully understand the advantages of cooperation in ecological communities (14) and to implement them in synthetic communities has been challenging (15–17). The focus has been on metabolic modelling to identify the keystone “currencies” of cooperation (9, 18–23). Knowing which metabolites are “traded” allows us to understand the mechanism of cooperation and how it enables survival in different environmental conditions. However, to emerge as the dominant strategy in any ecological niche, cooperative consortiums must grow faster than other free-living species. Metabolic modelling approaches can only determine the metabolic viability and synergy, in terms of the consortium’s productivity of metabolic flux exceeding the sum of individuals. Such models usually do not yield the growth kinetics of the cooperative consortium, and hence it has not been possible to study theoretically, the evolution of cooperation using existing evolutionary frameworks like fitness landscapes.

Furthermore, there are several shortcomings in the existing modelling approaches. Microbial growth is often posed as an optimal resource allocation problem that operates on the steady-state values of intracellular components, to simplify computation. Growth is estimated using the optimal steady-states values. However, often computation of the steady-state itself uses an estimate of the growth rate as the rate of dilution due to cell division, which leads to inaccurate estimates of growth. Often such models consider only a part of the cell and coarse-grain its dynamics to attempt maximisation of the energy generation while minimising the resource investment in catabolic enzymes and transporters (24, 25). More sophisticated whole-cell models like bacterial growth laws account for the shortcomings by computing a proxy measure of growth rate using the steady-state value of ribosome fraction of the proteome (26–29). Metabolic flux models entirely circumvent this by only computing the steady-state fluxes to optimise the genome-wide flux contribution to biomass and uncover the pathways or enzyme reactions that bottleneck growth (30–34).

These steady-state approaches, neglect the cell-to-cell differences due to stochastic gene transcription (35–37), leading to differences in the numbers of proteins (depending on the controlling promoter) (38), and in the cell’s metabolic state (39). While such variations are unimportant for prediction of the mean characteristics of the population, it is critical to model the growth of a cooperative consortium with cell-to-cell interactions such as the exchange of metabolites.

In this work, we develop a microbial growth model that not only captures the effects of stochastic variation on growth of non-interacting cells but also is applicable for studying the growth of cells that exchange metabolites. We implement stochastic burst-like gene transcription using Monte Carlo methods following the framework proposed by Golding *et al*. (40). To determine when a cell divides, we combined the concept of a biomass objective function (41) with the empirical laws for bacterial cell growth and size homeostasis: Adder and Sizer (42–45). While Adder proposes that cells divide upon addition of a fixed length to the birth size, Sizer proposes that division occurs when the cell grows to a characteristic length. We postulate that any cell size increment corresponds to an equivalent quantity and stoichiometric composition of biomass precursor metabolites. Thus, the cell divides only after producing the required quantity of metabolites in the correct stoichiometry.

Our framework attempts to capture the dynamics of cooperative growth and study its collective growth kinetics when cells exchange metabolites. We want to find the factors that lead to faster growth of the consortium, such that we can begin to investigate the evolutionary relationships between these. We obtained the distribution of cell doubling times, which mimics the experimentally observed distributions (46, 47) obtained from single-cell growth experiments run on microfluidic devices (42, 48). Our model successfully recapitulates the hyperbolic relationship between the substrate and growth rate observed by Monod (49). We demonstrate that gene expression noise manifests as noise in metabolite levels, which delays cell division (i.e. the noise is time additive). We also demonstrate that by reducing the effective number of limiting metabolites in a cell, either by auxotrophy or by the direct import of the metabolite, can accelerate growth. Lastly, we demonstrate that the growth rate of reciprocal cross-feeding auxotrophs can be greater than the prototrophs. In all, we demonstrate how incorporating stochastic gene expression noise into a cellular growth model allows us to capture the growth effect of auxotrophy and metabolite exchange directly, and how it can help understand the evolution of cooperation.

## Results

### 1 Model Development

We develop a cellular growth model that avoids relying on cellular steady-states and also incorporates the stochastic variation in growth.

As discussed, recent studies have put forth the empirical size based laws for bacterial cell division: Adder and Sizer (42–45). Combining these with the idea of biomass objective functions, we argue that cell size increase depends upon the production of equivalent quantity of constituent metabolites, in proper stoichiometry. We propose the idea as the “metabolic adder” and “metabolic sizer”, and use it as the foundation of our proposed cell growth model. Cell division is triggered only when the cell produces the minimum amounts of all the necessary metabolites (Fig – 1a). We demonstrate that our model approach recapitulates many of the known cellular phenomena and go on to extend the model to describe the kinetics of single-cell growth while they exchange metabolites among each other. We find the Adder and Sizer models very similar in their outcomes and proceed with Sizer for all the rest of the simulations in this work, primarily because it is found to be the better fit for the outcomes in case of poor media conditions, where growth in on minimal media with single substrates (44) (See Supplementary S4).

**Figure – 1.**
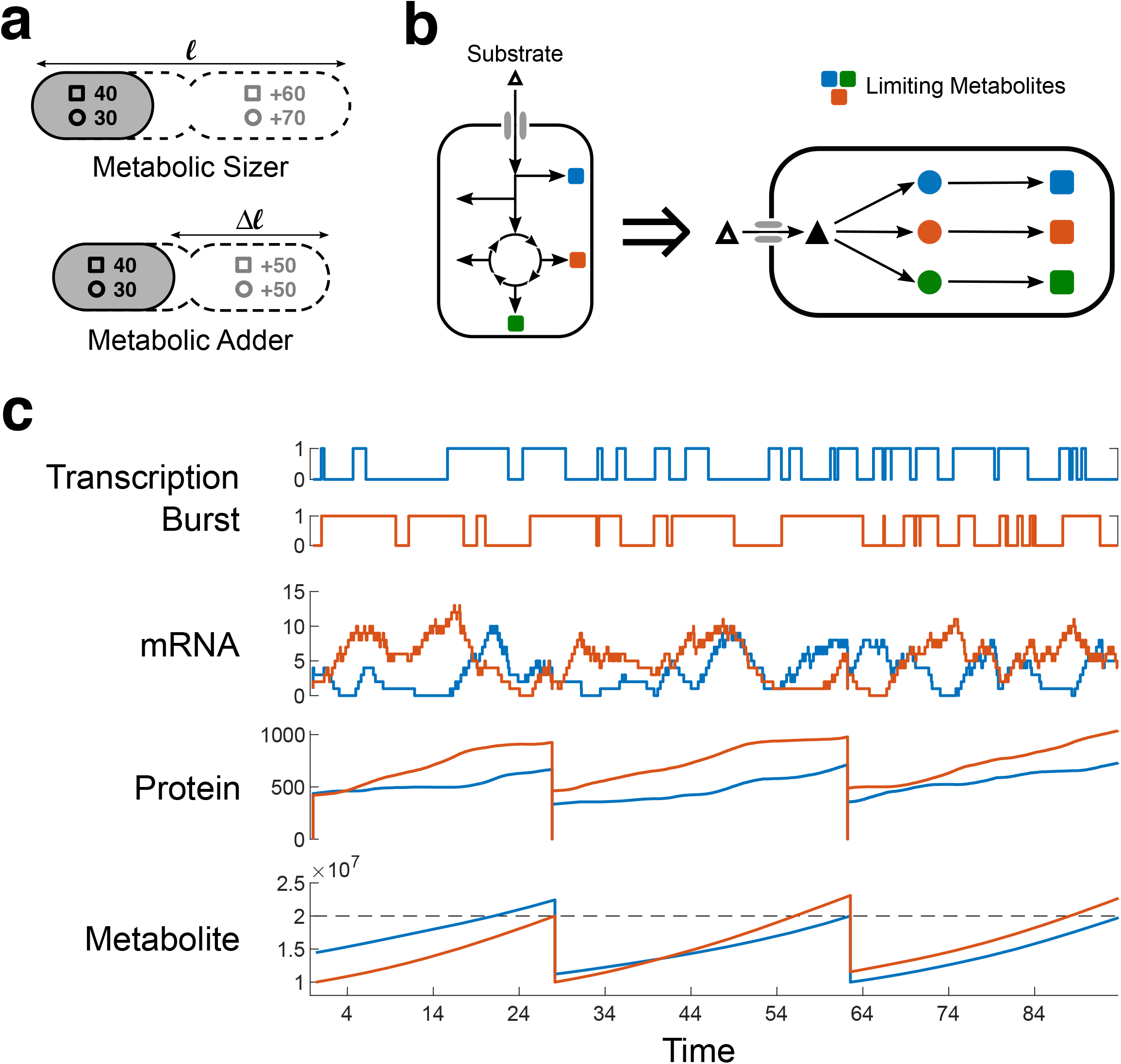
Model Schematic and behaviour of internal variables. (a) Model Schematic: Bacteria uptakes the substrate molecules from the environment and processes it via metabolic pathways to produce all the necessary metabolites for survival. Some (*p*) metabolites (coloured squares) bottleneck cell growth, since sufficient quantities cannot be produced in time. Our model assumes a simplified arrangement for all these limiting metabolite anabolic pathways, as linear enzyme cascades with *n* enzyme steps, all fed by a common substrate supplied at a constant rate. We consider all enzymes are equivalent in terms of expression and kinetic properties and are expressed in stochastic bursts. (b) Our model implements a “metabolite adder/sizer” model, where in we reinterpret the empirical size-based adder and sizer laws, in terms of the metabolites necessary for cell size increase. Such that cell division is triggered upon production of these metabolites in required amounts and stoichiometry. Always a fixed amount in case of Adder, and the balance amount in case of the Sizer. (c) Considering our model for *p* = 2 limiting metabolites and the metabolic Sizer law. The model implements stochastic burst like gene expression as a random process of transcription bursts switching on and off. During the burst, mRNA are produced, which act as template for protein translation. Both mRNA and proteins decay stochastically but mRNA decay much faster. The varying protein enzymes allow the production of the required metabolites at varying rates. According to Sizer, cell divides when both the metabolites cross the threshold (dotted line). Post division, cellular contents are halved.

Now, while theoretically, a shortfall in any metabolite could prevent a cell from dividing, only a few metabolites at a time may act as bottlenecks to cell division. These may be required in substantial amounts, or the production of the respective enzymes may be variable and in low numbers (39), such that the metabolite demand cannot be met in time. Additionally, some metabolites may be prone to leakage (may serve as public goods) and hence may act as a bottleneck to growth (50). We assume that the cell comprised of *p* anabolic pathways, whose products act as a bottleneck for cell division. All metabolite biosynthesis (anabolic) pathways are fed by the substrate supplied by the upstream catabolic pathways via different shunts from the Central Carbon Metabolism. We simplify the arrangement and assume that all anabolic pathways are fed a common substrate, which we further assume to be supplied at a constant rate *p. S_in_* per second (Fig – 1b).

The distribution of cell doubling times across different growth conditions collapse to one distribution when scaled by their mean (47). This leads us to postulate that the common downstream metabolic pathways like anabolism are responsible for the observed noise in growth, since in different growth conditions, upstream catabolic pathways may differ. Hence, our modelling approach focuses on quantifying the noise from anabolic pathways alone. Furthermore, to circumvent the differences due to the exact structure and kinetic properties of different anabolic pathways, we assume all *p* pathways to be linear and comprised of *n* enzyme catalysed steps, all of which we assume to be identical in kinetic properties and the transcription parameters.

We further assume that these *n* enzymes in each pathway are present on the same operon, and hence, the *n* enzymes share the same transcriptional burst profile (Fig – 1c). To incorporate the effect of stochasticity in our model, we implement stochastic burst-like gene expression, using the findings of exponentially distributed durations of transcription burst (*t_ON_*) and the waiting time between successive bursts (*t_OFF_*) (40). We randomly sample OFF and ON durations iteratively from exponential distributions of set mean, to generate the stochastic burst profile (Fig – 1c). Next, assuming no shortage of RNA polymerases, transcription proceeds at a constant rate till the end of *t_ON_*, where any incomplete transcripts are terminated. Each mRNA produced is assigned a lifetime by sampling from an exponential distribution. During the lifetime of an mRNA, one ribosome binds at a time and produces proteins, assuming ribosomes are always sufficient. (Any ribosome limitation only reduces the production of proteins and represent a nutrient-poor condition, but will not affect the overall dynamics). Any incomplete protein is terminated once the mRNA decays. Each protein produced is assigned a lifetime by sampling from an exponential distribution.

We use a system of coupled ODEs based on Michaelis-Menten kinetics for the enzyme kinetics, with the stochastic protein profiles as time-varying parameters to compute the production of metabolites (See Methods for the equations). We convert all quantities to molecules per cell assuming a cellular volume of *1μm^3^*.

All the biosynthetic pathways compete for the common substrate, and the flux they can acquire depends only on the number of enzymes present at each time, since kinetic parameters are assumed equal. When the production of all the *p* metabolites cross the set threshold based on the “metabolic adder or sizer” law used, the cell divides. Post division, all the contents of the cell are divided between the two daughter cells equally (51). The daughter cells start from the inherited values of the mRNA, proteins and metabolites, but generate unique stochastic burst profiles going forward.

### 2 Model properties and outcome

Simulating our model, for a given set of parameters, we obtain the duration of time between successive cell divisions or the cell generation time. The histogram of the generation times for all simulated parameter sets reveals a right-skewed distribution with a long tail, which concurs well with many experimental observations of bacterial growth at a single-cell resolution (42, 44, 45, 47, 52, 53). These studies indicated that the observed generation time distribution resembled a Gaussian on a logarithmic scale, and hence may be a Gamma or log-Normal distribution (47). Pugatch, on the other hand, analysed the single-cell dataset extensively and made a distinction between the quantities: cell generation time (*T_μ_*) (or inter-division time) and the cell doubling time (*T_div_*), the time for the doubling of cell size (46).

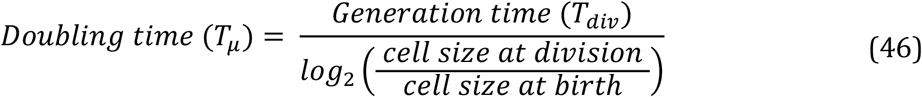

He tested the fit of different distributions to the experimentally derived doubling time distribution and found that a log-Fréchet or log-GEV (Type - II) distribution provided the best fit to the data.

Now, in our framework, cell division is triggered by exact size laws and division yields perfect halves, hence the ratio of size at division and birth is always two and thus generation time and doubling time are the same. We tested various combinations of gene expression and metabolite threshold parameters to obtain distributions that resemble physiology (See Supplement S3). The simulated generation times obtained using the selected parameters, give a long-tailed right-skewed distribution which is fit best by a log-Generalised Extreme Value (GEV) Type – III distribution (Shape parameter: k < 0) (Fig – 2a) (See Supplementary Fig – S3c, for comparison of GEV distribution shapes). Since the generation time distribution obtained is the maximum of the production times of the *p* metabolites, the obtained GEV-like distributions match with intuition derived from the Extreme Value theory (as the limit distribution of the maxima of a sequence of independent and identically distributed random variables) (54).

**Figure – 2.**
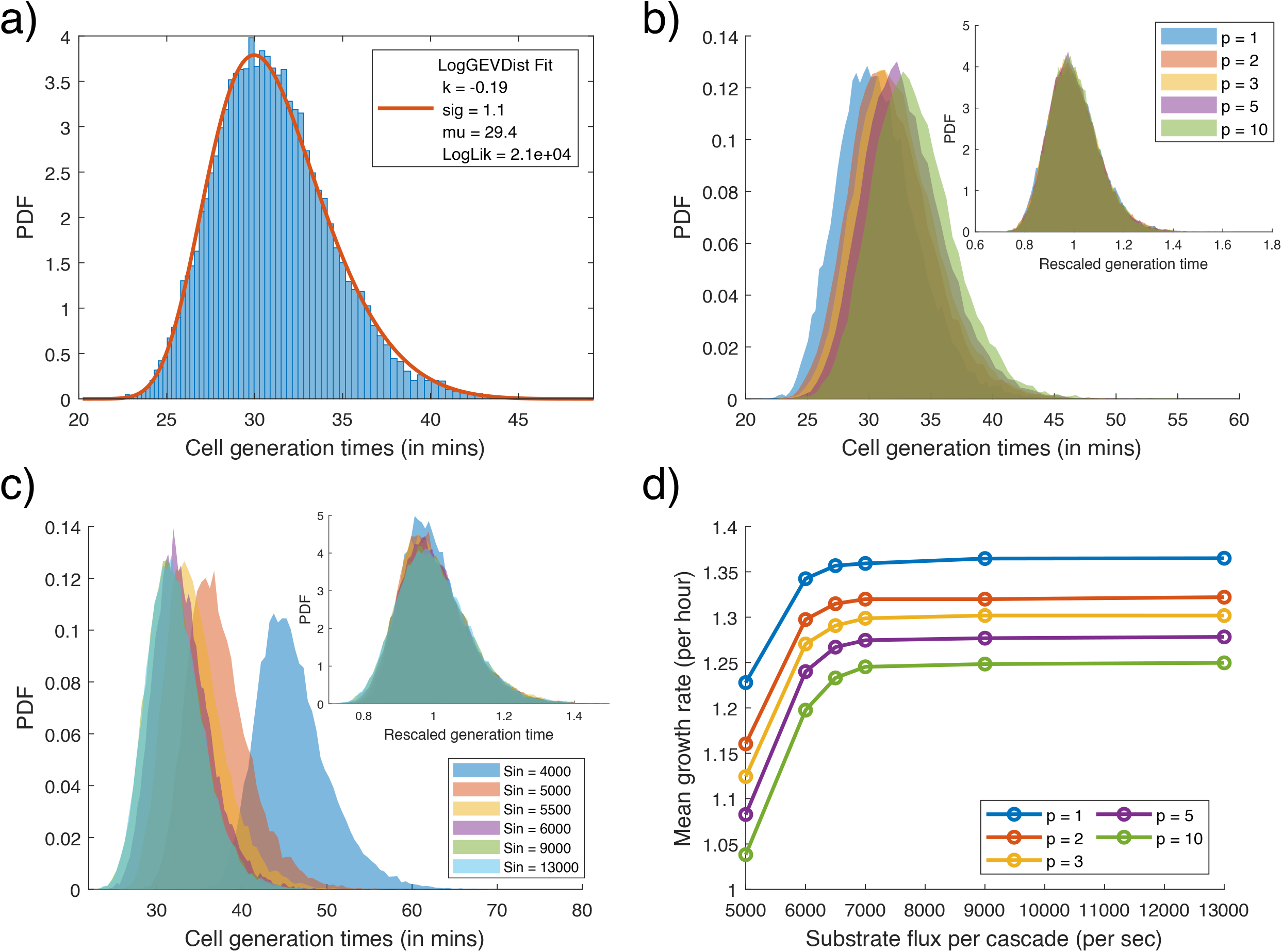
Model properties. (a) Distribution of cell Generation times obtained from model simulation is best fitted by log-GEV distributions. Shown distribution for simulation with *p* = 3, and *S_in_* = 9000/sec. (b) Comparison of distribution of generation times for varying number of limiting metabolites (*p*), at a high rate of substrate flux (*S_in_* = 9000/sec). Inset shows the distributions rescaled to their means. (c) Comparison of distribution of generation times varying the substrate flux rate (*S_in_*) in the simulations, for the case of p = 3 metabolite bottlenecks. Inset shows the distributions rescaled to their means. (d) Comparison of the mean growth rate of the simulated cells with varying number of bottleneck metabolites (*p*), for different rates of substrate flux (*S_in_*). The hyperbolic relationship is reminiscent of Monod’s Law.

Now, if the number of bottleneck pathways (p) increases, the chance that stochastic variations in enzymes would prevent the cell from reaching production thresholds at a given time, increases. Hence the maximum of the first passage times from *p* pathways, i.e. the generation time increases. The distribution of cell generation times, shifts to the right and reduces in skewness (Fig – 2b; Supplementary Fig – S1a). Moreover, upon scaling the data with the mean, we find that the distributions overlap (Fig – 2b inset), but with increasing *p*, the Coefficient of Variation (CV) of the distribution increases, while skewness and kurtosis decreases (Supplementary Fig – S1a).

Now, substrate flux per pathway (*S_in_*) represents the available concentration of the nutrients in the environment. The downstream anabolic pathways utilise the substrate for metabolite production. When *S_in_* is low, the rate of metabolite production is low and cell generation times are longer but as *S_in_* increases, metabolite production picks up, and cell generation times are shorter, and we observe the distribution of cell generation times shift to the left (Fig– 2c). Upon scaling the data with the mean, the distributions overlap (Fig – 2c inset), but with increasing *S_in_*, skewness and kurtosis decrease, and distribution CV also increases and saturates (Supplementary Fig – S1b). Next, when we plot the associated population growth rates against the corresponding *S_in_* values, we observe a hyperbolic relationship, for all values of *p* simulated, which recapitulates Monod’s observation of a hyperbolic relationship between substrate concentration and the cell’s growth rate (49) (Fig – 2d).

Substrate flux is known to be the primary determinant of growth rate in a cell, and studies have hence focused on quantifying the noise in catabolic pathways as a measure of substrate flux variation to estimate growth rate variations (36). However, during balanced growth conditions, the substrate flux from catabolism is stable, and hence we chose only to model the noise contribution from the downstream anabolic processes.

Studies have shown that the cell division control is controlled by more than one factor and limiting agent (55, 56), which in our model is represented by multiple bottleneck pathways (*p*). In a real cell however, the number of bottlenecks may depend on the cellular growth conditions, and in case of poor media conditions (low *S_in_*) it may increase. Although our model does not have the ability to perform such internal metabolic switches, it nonetheless can still recapitulate physiological observations such as the distribution of generation times and Monod’s Law (Fig – 2d).

### 3 Effect of metabolite uptake and secretion on simulated cells

Our primary goal is to develop a framework that can quantify the effect of nutrient uptake and secretion on the growth of cells. Our model design captures the effect of importing external metabolites and exporting internally produced metabolites, directly in terms of the generation time.

Till now, our simulations have considered cells were grown on media that provides only the substrate flux (*S_in_*). We now simulate growth when additional limiting metabolites are present in the media (like amino acids, nucleotides). Our model cell imports these metabolites directly at a constant rate to meet the internal metabolite requirement. The import effectively lowers the metabolite’s threshold quantity that needs to be produced internally and thus, cells on average divide faster. Since we do not explicitly model any metabolite feedback, the substrate flux availed by the biosynthetic pathway does not reduce, and the substrate competition remains same (We relax this condition later in Results Section 6). With increasing import rates of the limiting metabolite, the generation times shift to the left (Fig – 3a and 3b), and the growth rate increases (Fig – 3c). From our model’s perspective, importing a limiting metabolite at a high rate makes it non-limiting. Thus, decreasing the number of bottlenecks (*p* → *p* — 1) removes its noise contribution to cell division (Fig – 4d), allowing the cell to divide faster. The growth advantage from the import of a limiting metabolite has an upper bound, and the advantage from a higher rate of import saturates quickly (Fig – 3d).

**Figure – 3.**
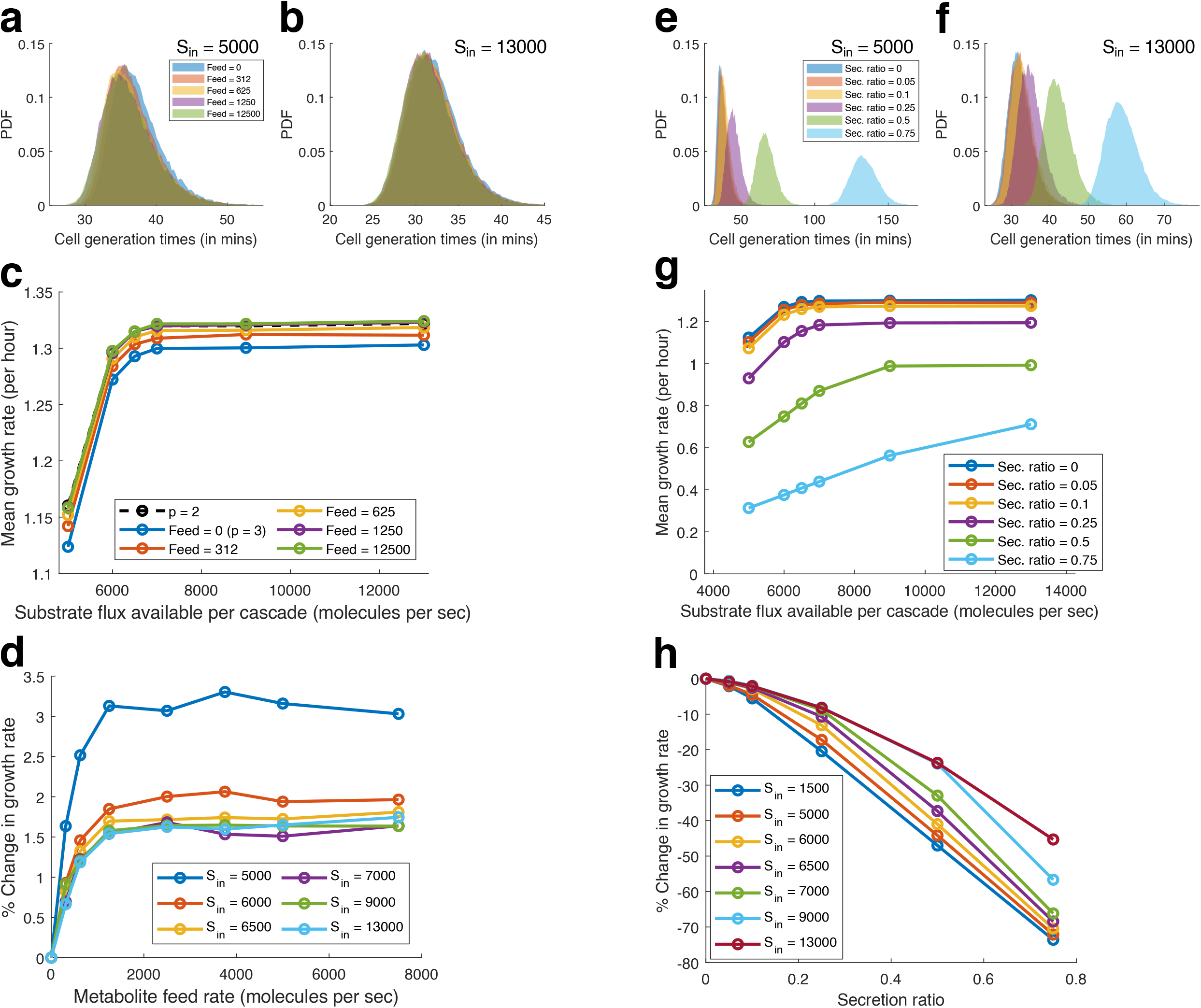
Effect of metabolite uptake and secretion on simulated cells. (a-d) **Effect of metabolite uptake**. Comparison of generation times for different rates of metabolite import, (a) shows low substrate flux and (b) shows high substrate flux. (c) Comparison of the mean growth rate of the simulated cells with varying rates of metabolite import, for different rates of substrate flux (*S_in_*). (d) Comparison of the relative change in growth rate compared to the case of no uptake, due to varying metabolite feed (uptake) rates, for different substrate flux (*S_in_*). (e-h) **Effect of metabolite secretion** Comparison of generation times for different ratios of metabolite secretion (fraction of produced metabolite secreted), (e) shows low substrate flux and (f) shows high substrate flux. (g) Comparison of the mean growth rate of the simulated cells with varying metabolite secretion ratios, for different rates of substrate flux (*S_in_*). (h) Comparison of the relative change in growth rate compared to the case of no secretion, due to varying metabolite secretion ratios, for different substrate flux (*S_in_*).

**Figure – 4.**
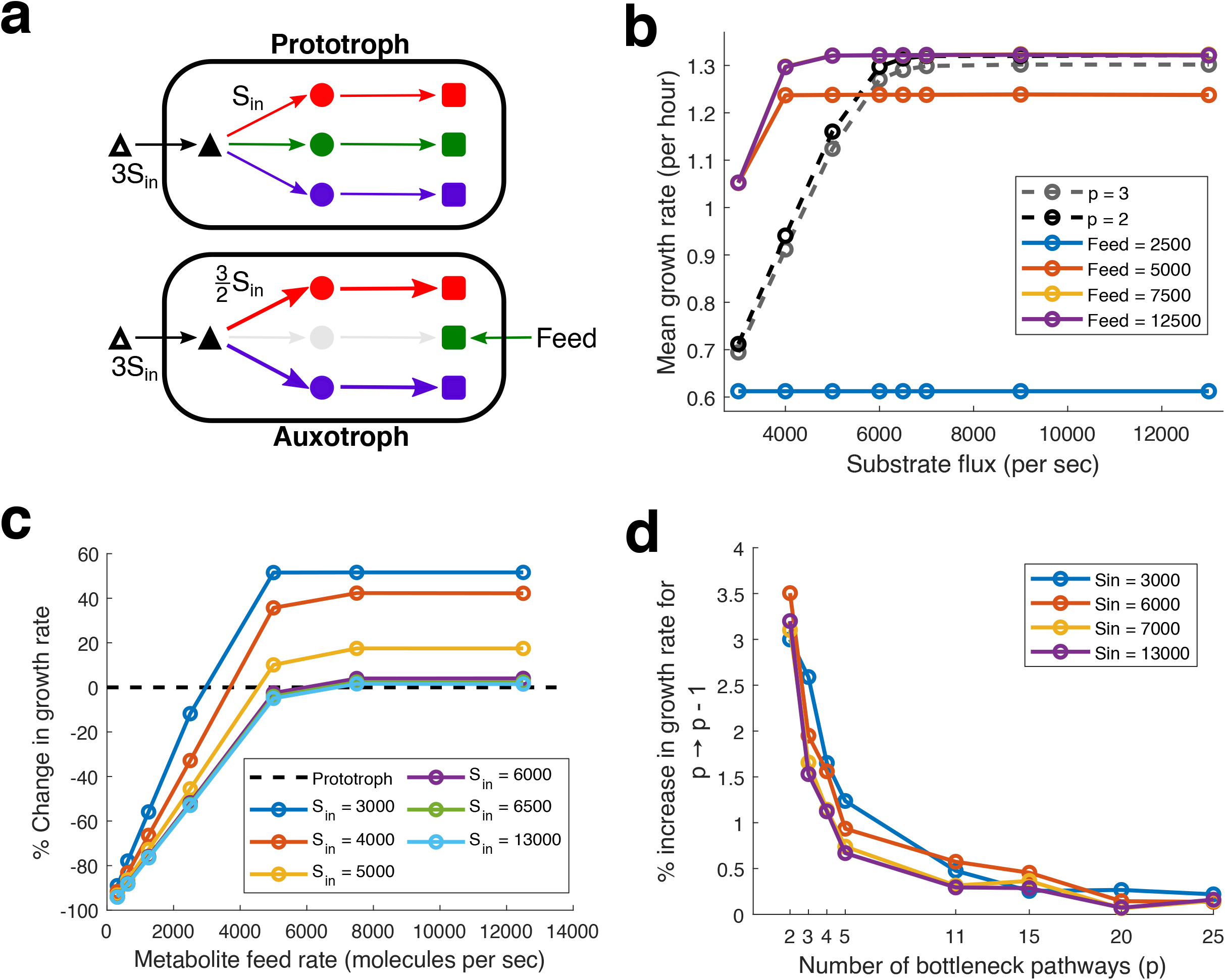
Growth advantage of auxotrophy. (a) Schematic figure comparing an auxotroph (*p =* 2) with the ancestral prototroph (P = 3). (b) Comparison of the mean growth rate of the simulated auxotroph with varying metabolite feed (uptake) rates for the missing metabolite, for different rates of substrate import flux (*S_in_*). The grey and black dotted lines represent the prototrophs with *p =* 3 and *p =* 2 bottleneck metabolites respectively. (c) Comparison of the relative change in the growth rate of the auxotrophs, when compared with the original prototroph (*p =* 3), for varying rates of metabolite feed, and substrate import flux (*S_in_*). (d) Comparison of the percentage increase in the growth rate upon loss of a bottleneck pathway for different starting number of bottleneck pathways (p), simulated at different substrate import flux rates (*S_in_*).

Next, we consider the effect of exporting metabolites from the cell. Some metabolites are usually produced in excess and leak out from the cell (10–12, 50), without any adverse effect on growth. However, if these metabolites are not useless by-products for the cell, then excess secretion could impede growth. In case, one of the limiting metabolites is secreted, it leads to an effective increase in its metabolite threshold, and hence the cell requires more time to divide. We quantify the growth effect of secreting metabolites, by simulating secretion as a fraction of the total metabolite produced per unit time, rather than a constant amount secreted per unit time, since it synchronises the secretion to the internal production. It hence captures an accurate estimate of the amount secreted (low secretion after birth due to low enzyme, and higher secretion before division due to a higher number of enzymes and transporters). As the secretion ratio increases, cell generation times increase, and the distribution shifts to the right (Fig – 3e and 3f). The magnitude of the difference is however much more pronounced in case of a nutrient-poor media (low *S_in_*, Fig – 3e). With the distributions scaled to the mean, we further see that increased secretion leads to lowered skewness, and skewness decreases more in case of the nutrient-poor media (Supplementary Fig – S1d). Unlike the case with nutrient uptake, the effect of secretion is unbounded, and increased secretion further lowers growth rate (Fig – 3g and 3h).

### 4 Growth advantage of auxotrophy

We have already demonstrated how direct import of a limiting metabolite can increase growth, and how the advantage is capped. We described this phenomenon as the partial alleviation of the noise contribution from one of the bottleneck metabolites, which corresponds to (*p* → *p* — 1) in our model. If the modelled prototrophic cell mutates to lose one of the biosynthetic pathways for a limiting metabolite (and becomes an auxotroph), then the same outcome may be obtained. Interestingly, this is a common phenomenon, seen not only in endosymbiotic bacteria (57) but also in some free-living bacteria (58, 59). Black Queen Hypothesis proposes that this genome reduction is driven by a resulting selective advantage (50, 60). However, such gene loss makes the survival of the cell contingent on sufficient external supply of the metabolite.

For the simulation, we consider a cell with three bottleneck pathways (p), and then delete one of them (Fig – 4a). The cell directly imports the metabolite it cannot synthesize. Additionally, the available internal substrate flux (p. *S_in_*) originally meant for p pathways, is now redistributed among the remaining p — 1 pathways. The auxotroph originally with three bottleneck pathways, now receives 50% more substrate flux in the two remaining bottleneck pathways and hence divide faster (Fig – 4a). At low import (Feed) rates, this missing metabolite determines the cell generation time, however at higher import rates the missing metabolite is no longer the bottleneck, and the generation time distributions overlap with the distribution for p — 1 (i.e. p = 2) bottlenecks (Supplement S8), as expected. It is interesting to note that when an auxotroph is grown in poor media conditions, represented by the low substrate flux (*S_in_*) in our simulations, a sufficient rate of import of the limiting metabolite, allows the cells to grow much faster (Fig – 4b and 4c), relative to the growth advantage observed for high substrate flux conditions.

Our model does not incorporate the effect of saved protein costs, however the effect of saved substrate flux is incorporated and reflected in the growth advantage at low substrate flux (*S_in_*). Moreover, even when the effect of substrate flux saturates, the model predicts a growth difference between the ancestor prototroph and the auxotroph. Our model presents a novel factor contributing to the growth advantage due to gene loss, in addition to resource savings (in terms of protein cost and substrate flux) (50, 60), leading to the spontaneous evolution of auxotrophs (61).

It is imperative to note that the magnitude of growth advantage demonstrated here, corresponds to a low number of bottlenecks (low p). If a large number of metabolites are bottlenecks simultaneously (large p), then the relative growth advantage observed due to loss of one pathway or direct import of one limiting metabolite, will be significantly lower (Fig – 4d). While it is known that for different nutrient media, different metabolites act as bottlenecks for growth (62), it is not immediately clear which biomass precursor metabolites act as a bottleneck, how their numbers vary, and their relative limiting effect on growth. In our model, we considered all bottlenecks of equal strength, for simplicity.

### 5 Towards reciprocal metabolic cross-feeding

As discussed, the survival of auxotrophs is contingent on the external availability of the metabolite they cannot produce internally. These cells invade ecological niches where other cells secrete the necessary metabolites, either as by-products or public goods. In a public-goods scenario, the invading cells act as “cheaters” and reduce community productivity (63–65). However, if the metabolite is a by-product and its accumulation toxic, then its clearance from the environment could induce the producer cells to secrete more, and improve community productivity (8, 66, 67).

Analysis of available microbial genomes from various niches has revealed that most of them are auxotrophic for at least a few essential metabolites, such as amino acids (60, 68). Although surprising, it explains why ecological examples of metabolite cross-feeding are so commonplace (1–5). During the process of metabolite cross-feeding, the metabolites secreted by the microbes diversify the nutrient composition of the environment, which enables survival and further evolutionary diversification of the group, by allowing for spontaneous evolution of auxotrophs (61).

However, prototrophic species could invade such cross-feeding populations, and by importing the secreted metabolites “cheat” to grow faster and outcompete the consortium (69). To avoid such a fate, cross-feeding must also enable fast growth, enough to surpass the competing species, in addition to enabling the survival of the group. Nevertheless, we do not understand how the division of labour may allow cooperating cells to improve productivity enough to exhibit growth that is faster than prototrophs.

We have demonstrated that the import of limiting metabolites accelerates growth, but on the other hand, secretion of such limiting metabolites at high rates can also slow down growth. In a cross-feeding interaction, cells reciprocally exchange metabolites, by secreting one metabolite and importing another simultaneously. How must cellular resources be balanced between these processes to achieve a net positive outcome is not understood, even for a simple symmetric exchange.

Let us consider two reciprocal auxotrophs, each overexpressing the metabolite the other needs for survival. These auxotrophs due to loss of one of the metabolic biosynthesis pathways, grow faster because they can redirect the saved resources from the pathway towards growth, and also reduce the effect of gene expression noise on growth (as we demonstrate using our model). However, the need to supply metabolites to each other requires the cells to invest the saved resources towards metabolite overproduction, taking away the growth advantage. Assuming, metabolites of similar costs are exchanged, intuitively, we may expect that the reinvestment of saved resources may enable a doubling of the metabolite production, and hence allow the cells to exhibit a growth rate as high as an equivalent prototroph. Which is precisely the result we obtain when we simulate a simplified steady-state version of our model (Supplementary Fig – S5) (See Supplement S5 for details). That too, when not considering the possible losses and delays due to transport of the metabolites. Thus, it is hard to imagine that cross-feeding can enable such growth advantages.

Pande *et al*. used synthetic reciprocal amino acid auxotrophs that also overproduced amino acids to study this phenomenon, and found a large number of complementary auxotrophs when co-cultured could cross-feed stably, exhibiting growth rates faster than the parental prototroph in monoculture (11). However, only a few pairs could outcompete the prototroph in a competitive co-culture, since the prototroph could feed on the secreted metabolites. Here, we use our stochastic growth framework in an attempt to explain these observations.

We start with an auxotroph (*p* = 2) originally a prototroph (p = 3) (Fig – 5a). Next, we need to implement over-expression of the secreted metabolite (using the resources saved). Since in our model, pathway enzymes are equivalent, loss of a pathway allows another pathway to produce double the number of enzymes at maximum. However, it is unclear how a real cell handles the over-expression of a pathway. In our stochastic framework, over-expression involves changing the transcription burst frequency, but since it would be hard to estimate burst parameters that precisely “double” the enzyme, we instead double the kinetic constant of the enzyme keeping the same stochastic burst profile, to set up a fair comparison. We considered secretion at different ratios of the amount of metabolite produced per unit time. We further varied the direct import rate of the missing metabolite. We study the growth of this auxotroph that produces twice the quantity of metabolites per unit time, while importing the missing metabolite, as a proxy for growth of cross-feeders (Fig – 5a).

**Figure – 5.**
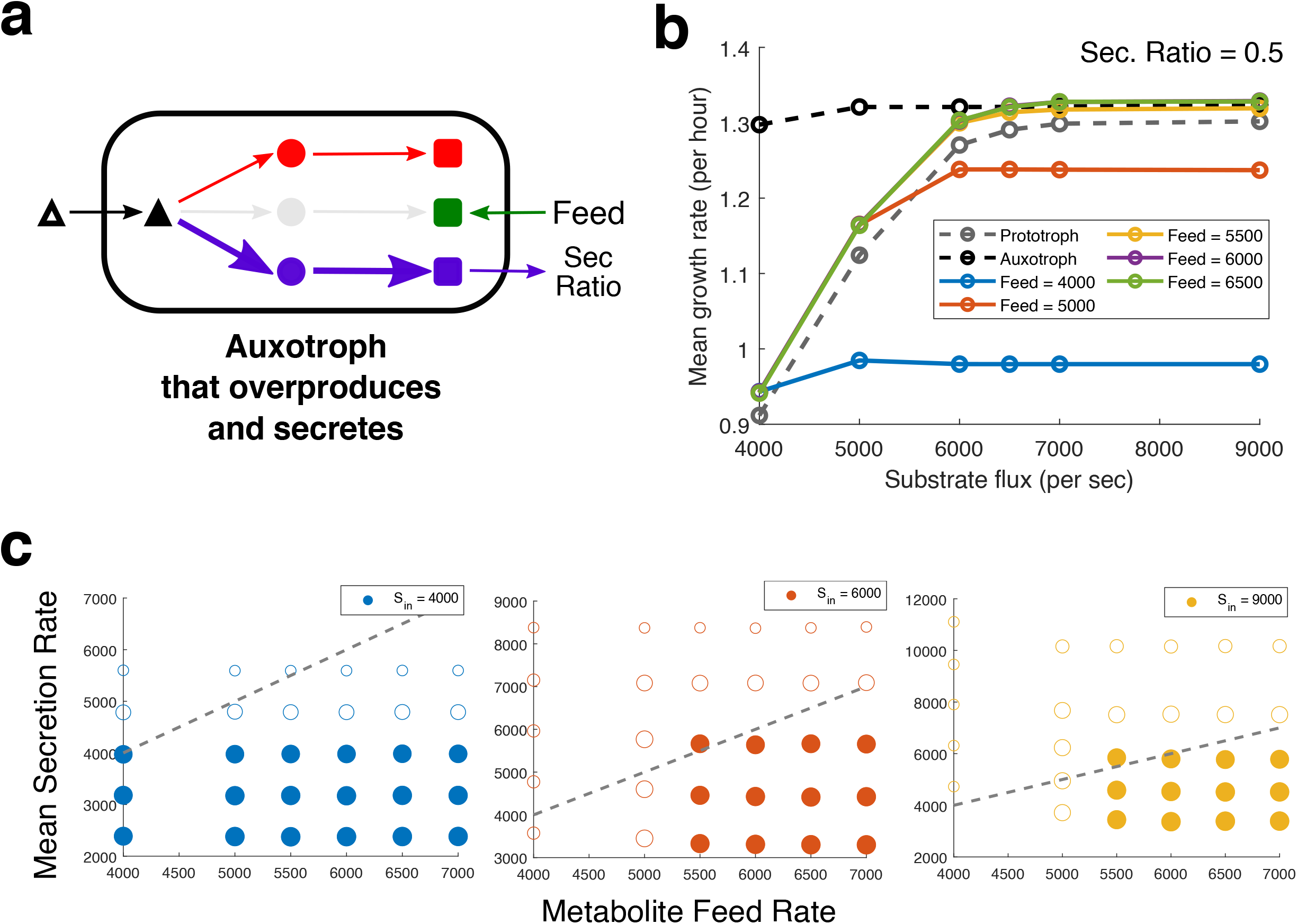
Reciprocal metabolic cross-feeding. (a) Schematic figure depicting our proxy for reciprocal metabolic cross-feeding: an auxotroph that overexpresses and secretes one the limiting metabolites (Blue), while importing the metabolite it is unable to produce (Green). (b) Comparison of the mean growth rate versus different rates of substrate import flux (*S_in_*), for auxotrophs that overexpress (2X) and secrete (0.5X) a limiting metabolite (proxy for cross-feeders). The grey and black dotted lines represent the growth profile of corresponding prototroph and auxotroph. (c) Comparison of the mean secretion rate of metabolites (Y), for an input metabolite feed rate (X), for different substrate flux rates (*S_in_*), represented as different colours of the plotted circles. The size of the circles represents the magnitude of the growth rate. The circles are filled if the simulated cells grow faster than the prototroph (*p =* 3) when grown without any additional metabolite feed. The dotted diagonal line represents equality of secretion and feed rates.

We find that the proxy cross-feeders demonstrate growth rates higher than the prototroph, even when they secrete 50% of the metabolite they overproduce (Fig – 5b). For these overproducing and secreting auxotrophs to sustain a feasible cross-feeding interaction, they need to secrete metabolites at a rate which is higher or equal to the rate at which they import metabolites. We plot the mean secretion rates for each simulated metabolite feed (import) rate at different values of substrate import rate (*S_in_*) (Fig – 5c). The dotted lines represent secretion rate is equal to the metabolite feed. Thus, points above the diagonal represent secretion rates is higher than the metabolite import. The radius of the circles represents the growth rate of the simulated cells, and the circle is filled if the growth rate is higher than the growth rate of the ancestor prototroph (without any metabolite feed). Thus, filled circles above the diagonal represent “feasible” parameters where cross-feeding cells can exhibit faster growth while maintaining higher overall metabolic productivity. In Fig – 5c we find such feasible solutions at mid and high values of *S_in_*. Thus, kinetic advantage of cross-feeding consortiums may only be observed when sufficiently concentrated substrate is available.

We demonstrate that even in the case with minimal headroom for improvement of growth rate (i.e. when the competing bottleneck pathways are of an equal threshold), feasible solutions for cross-feeding can be found. In cases where the subsequent bottlenecks are of a lower threshold, the cell would have a higher headroom to improve its growth with higher import rates. However, lower growth durations may also lead to lower secretion. In a real cell, however, faster growth also implies faster DNA replication, which would then allow higher protein production, which would lead to higher metabolite production.

### 6 Effect of feedback regulation on metabolite biosynthesis and the effect of noisy metabolite import

#### Effect of feedback regulation

When the metabolites synthesized in the cell are available in the environment, cells readily import them into the cell, inhibiting the internal biosynthesis due to feedback regulation. The substrate flux utilized by the pathway is reduced, and more substrate flux is available for use by the other competing biosynthesis pathways. Till now, in our simulations, we have neglected this effect. Here we implement the inhibition due to feedback regulation into our model by incorporating a scaling parameter multiplied with the enzyme’s catalytic constant, which reduces the effective flux through the pathway, making the metabolite imported directly. Thus, the competing biosynthetic pathways can utilize the excess substrate flux for production of other limiting metabolites.

By varying the scaling parameter, we vary the amount of feedback as a percentage reduction in the enzyme’s catalytic constant. We also vary the rate of metabolite import (Feed) into the cell and the substrate flux (*S_in_*) as done before (Results Section 3: Effect of metabolite uptake and secretion on simulated cells). At low metabolite import rates, increasing feedback strength leads to a decrease in the growth rate, since the metabolite production rate slows down (Fig – 6a). At high metabolite import rates, we find that increasing feedback strength makes more substrate flux available to the competing biosynthetic pathways, and hence the growth rate increases in low substrate flux (*S_in_*) cases (Fig – 6b). No further increase in growth rate is possible at higher substrate flux because the enzyme is limiting.

**Figure – 6.**
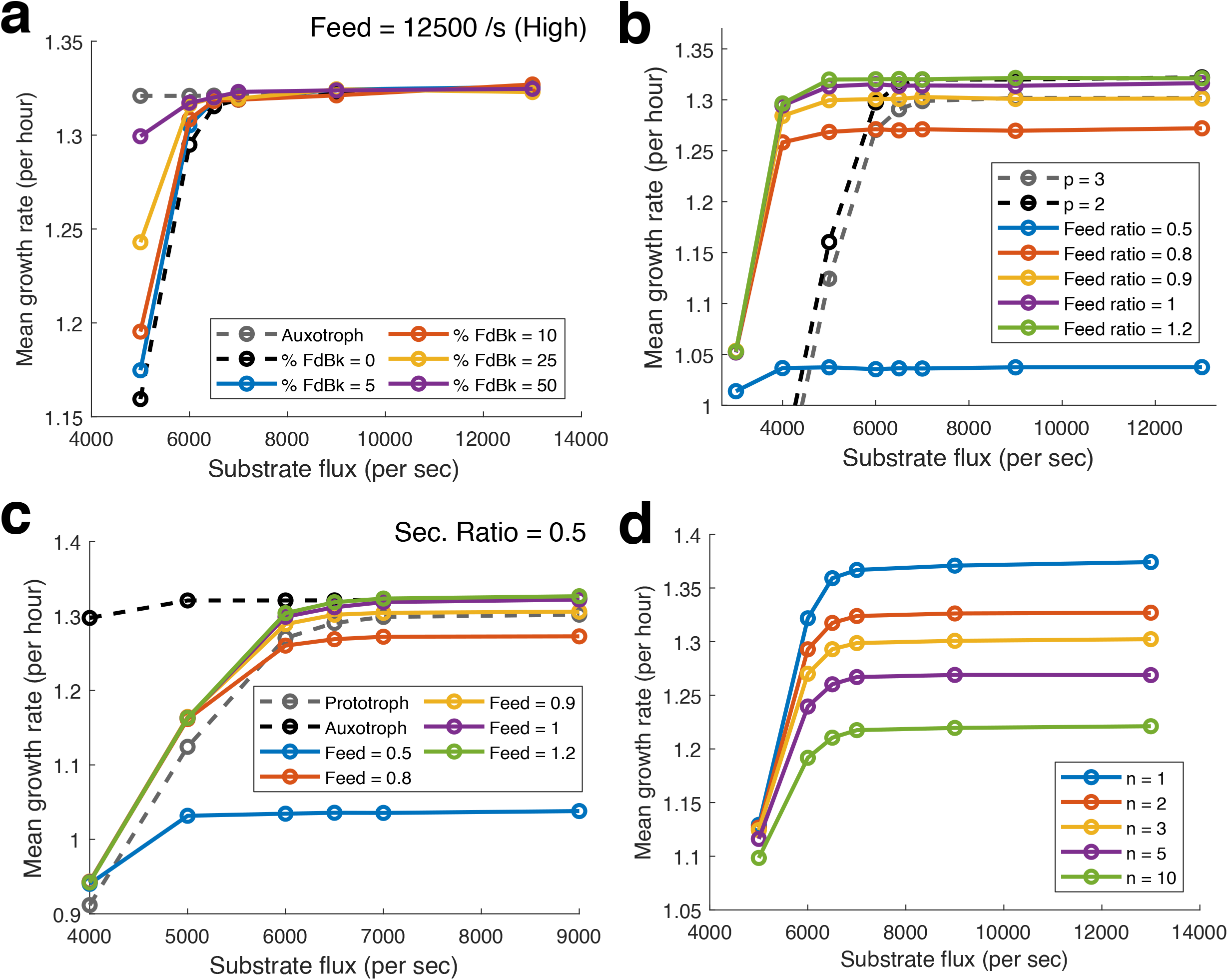
Effect of feedback on metabolite biosynthesis and noise during metabolite import. (a) **Effect of feedback on metabolite biosynthesis** Comparison of the mean growth rate of the simulated prototroph versus different rates of substrate import flux (*S_in_*), when fed a limiting metabolite at a high rate = 12500/s, with varying strengths of feedback regulation (%). Growth rate of prototroph with no feedback effects (black dotted line) and auxotroph (grey dotted line). (b-c) **Effect of noise during metabolite import** Comparison of the mean growth rate versus different rates of substrate import flux (*S_in_*), for (c) auxotrophs, and (d) auxotrophs that overexpress and secrete a limiting metabolite (proxy for cross-feeders). In (c) the grey and black dotted lines represent the prototrophs with *p* = 3 and *p* = 2 bottleneck metabolites respectively. In (d) the grey and black dotted lines represent the growth profile of corresponding prototroph and auxotroph. (d) **Effect of varying the number of enzyme steps in the biosynthetic pathway** Comparison of the mean growth rate versus different rates of substrate import flux (*S_in_*), for a prototroph with 3 limiting metabolites (*p =* 3), while varying the number of enzyme steps in the biosynthetic pathways (*n*).

#### Effect of noisy metabolite import

In our model, we assume that biosynthetic enzyme gene transcription is a stochastic burst-like process. However, this is true for all genes transcribed in the cell. Thus, the number of transporter proteins in the cell should also vary. Nevertheless, in our simulations, we assumed that metabolite import at a constant rate for the sake of simplicity.

Here, we explore how our observations may change when we model metabolite transport as a noisy process. We assume that the transporter protein’s expression parameters are the same as the biosynthetic enzymes, for ease of comparison due to identical noise characteristics. We also assume the same catalytic constant for the transporter as the enzyme. However, we use a scaling parameter (Feed ratio) multiplied to the catalytic constant to vary the rate of import.

We have shown that importing one of the limiting metabolites into the cell at a sufficiently high rate allows prototrophic cells to enjoy a growth advantage (Fig – 3c). We have also demonstrated that auxotrophs require a sufficiently high supply of the metabolite they cannot produce to enjoy a growth advantage (Fig – 4b). In the case of auxotrophs and cross-feeders, one may intuit that if the growth advantage is a consequence of reduced noise, then incorporating noise in transport may cause the observed growth advantage to disappear. We thus incorporate noise into the process of transport and then simulate their growth, as done before in Results Section 4 and 5.

Upon simulating the growth of auxotrophs with noisy metabolite import, we see that the growth advantage observed sustains, even when the import and biosynthesis have the same catalytic constant (Feed ratio = 1, Fig – 6b).

In the case of cross-feeding, where we study the growth of a metabolite overexpressing and secreting auxotroph as a proxy, we again observe that the growth advantage sustains with noisy metabolite import (Fig – 6c).

An important fact when comparing the noise due to biosynthesis versus the transport, is that biosynthesis involves multiple steps, while transport is always a one-step process. Our model considers that each biosynthetic pathway is linear and composed of *n* enzyme steps. Through this work, we have assumed 3 enzyme steps (*n* = 3). In Fig – 6d, we plot the growth rate of cells against the rate of substrate import, and vary the number of enzyme steps (*n*). We observe that the growth rate decreases as the number of enzyme steps (*n*) increase. Thus, even when transport is modelled as a stochastic process, the growth advantage sustains due to the difference in the number of linear steps between the processes.

## Discussion

We demonstrated that accounting for stochastic burst-like gene expression (40) captures variations in the production rates of essential cellular metabolites (biomass precursors) (39) and hence differences in cell growth and division kinetics. The framework applies to both non-interacting cells, as well as cells that exchange metabolites. Our model recapitulates fundamental relationships like Monod’s law and also yields the experimentally observed right-skewed long-tailed distribution of cell doubling times (Fig – 2). Furthermore, our framework can demonstrate the growth advantage of metabolite import (Fig – 3c) and genome reduction (auxotrophy) (Fig – 4b), without invoking the idea of protein cost savings, simply as an emergent consequence of the stochastic description, under the assumption of a strict metabolic Adder or Sizer law. We also show how a symmetric reciprocal cross-feeding system, which is expected to not enjoy any of the benefits of resource saving, may still enjoy a growth advantage over the ancestor prototroph (Fig – 5b). Lastly, we test how feedback regulation affects the outcomes, and demonstrate that the growth advantage persists even if we incorporate stochastic kinetics for metabolite import (Fig – 6).

Stochasticity in a bacterial cell growth model has also been used by Thomas *et al*. Int his work, the authors focus on quantifying the relative contributions of different sources of stochastic noise (such as transcription, cell partitioning, mRNA degradation, translation and other processes) leading to variations in the growth rate, across a wide range of mean growth rates (70). The work builds on their previous steady-state model (26), and used the Cooper–Helmstetter model (44, 71) with Donachie’s constant initiation cell size per origin (72) to determine when cells divide. The model focuses on growth rate variations which diminish at high rates, unlike the variation in cell generation times that persists even in high growth rates (42). The model does not connect the stochastic production of individual biomass precursor metabolites and cellular growth, and hence is unable to account for changes in growth upon their import or secretion.

As discussed in results Section 5, our model aims to be useful by directly yielding growth differences between different cellular states that arise in the ecological niche as microbes compete and cooperate. The participating microbes may be prototrophs, or auxotrophs. Additionally, individual cells may not only import metabolites but secrete them as well, at varying rates, controlled by internal expression and transport parameters. The framework we developed can determine the cell generation times associated with these cellular states and how it will change due to metabolic interactions, such as the case of reciprocal cross-feeding interactions between auxotrophs. Each of these represents intermediate states towards the evolution of cooperation, and this framework offers an opportunity to study if such an evolution towards cooperation is possible. Some studies have pitted genome-scale dynamic flux models of microbes with different auxotrophies and metabolite secretion rates (or leakiness) against each other (73, 74), some have even used an evolutionary game theory framework to derive intuition about which microbial combinations could be evolutionarily successful (75, 76). However, these models cannot be used to study the real-time emergence and fixation of such cooperative pairs, starting from prototrophs. Our framework on the other hand brings to life such a possibility, as it allows us to simulate the stochastic growth of individual cells in a population, as they interact metabolically. Our model however cannot accurately capture the dynamics of growth when the media conditions shift drastically, since our model cannot dynamically update its internal metabolite thresholds. (Adder and Sizer lengths increase with increased mean growth rates (42)). When limited to a constant media condition, combined with a mutational framework, we expect our model will help uncover new insights towards the evolution of cooperation, by identifying the environmental regimes where it could happen.

## Methods

### 1 Model setup and calibration

Our model ascribes no specific identity to the enzymes in the cell, and thus we assume the simulated enzymes to have average properties of all cellular enzymes. We obtain the average length of bacterial proteins, the median values of enzyme catalytic constant and enzyme half-saturation constant (See S9: List of Parameters). For the stochastic gene expression, we follow the scheme described by Golding *et al*. 2005. We estimate parameters and the associated metabolite threshold by simulating various combination of values, and select for mean generation time and the coefficient of variation (CV) closest to physiological values (See Supplement S3). We finally chose a metabolite threshold 1E7 for Adder, and 2E7 for Sizer, while *t_ON_* and *t_OFF_* chosen were 4 mins and 2.4 mins respectively.

### 2 Metabolic Adder and Metabolic Sizer

By using the stoichiometric quantity of metabolites produced as a proxy for the cell size, we obtain the metabolic adder and sizer models. For simplicity we assume 1E7 molecules of each limiting metabolite is required to make a new cell. Thus, in case of metabolic adder, 1E7 more molecules of each limiting metabolite must be produced irrespective of the inherited metabolites. In case of metabolic sizer, the cell’s total production of each limiting metabolite needs to exceed 2E7 to trigger cell division. Fig – 1a, represents this schematically.

### 3 System of coupled differential equations used to compute metabolite production

After generating the stochastic enzyme profile, we pass it as a time-varying parameter to a system of coupled ODEs, based on Michaelis-Menten kinetics. While it is possible to also consider the noise from the stochastic metabolic reaction processes, we disregard it in our model. The nature of this noise in effect only leads to fluctuations in the mean catalytic constant of the enzyme, which becomes important at the timescales of single-molecule experiments (77), and averages out and when we consider the metabolite production at the time scales of cell growth and division.

The primary substrate S common to all the pathways is supplied at a constant rate *p. S_in_* and each of the enzymes feeds off it. *i* represents the bottleneck pathways 1 to *p*.

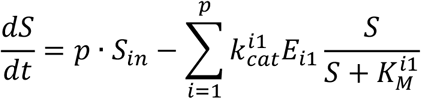

The subsequent products except the final products are modelled as:

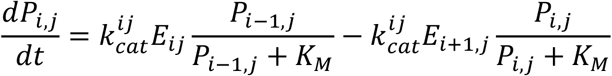

*j* represents the linear enzyme steps in each pathway.

The final step of the biosynthetic pathway is modelled as:

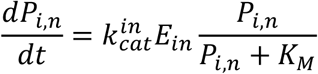

For our simulations, we considered each of the enzymes in the concurrent pathways equivalent in terms of their expression and kinetic parameters. Thus, 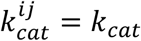 for all *i, j*.

### 4 Model Simulation

We start the simulation of the model with the obtained parameters, however for the lack of knowledge of any initial conditions we start with all zero. As we start the simulation, firstly stochastic mRNA and protein profiles are generated based upon the set gene expression parameters, which are then used as time-dependent parameters while solving the system of coupled ODE numerically using *ode15s* in MATLAB. When all the *p* metabolites have met their production requirements, the events function is triggered, which terminates the evaluation of the ODE, and marks the cell division. The difference between the end and start times of the ODE solution gives us the growth duration or generation time.

Before we start recording values from the simulation, we allow the system to saturate by letting the cell run through 100 divisions or generations, following only one of the daughter cells, starting from the zero initial values. Thereafter, we simulate an exponential population growth, by simulation all the daughter cells, till 13 generations, starting from 1 cell, and hence obtaining growth data for 2^13^ — 1 = 8191 cells. We use the generation times to compute the growth rate of the population, as described in the next section. We repeat the simulations with 3 or 5 unique seed values to obtain concordant observations.

### 5 Computing growth rate from generation times

From our simulations, we obtain the birth and division time of each simulated cell. If we sort the cells in the ascending order of their birth and consider each birth event as a unit increment to the population, it gives us the microbial population growth curve. The points appear as a straight line in a semi-log plot, however the last part appears to saturate because there are no more cell divisions. To estimate the growth rate, we consider only the first half of the sorted dataset, to obtain the log or exponential growth phase. We plot the best fit line on the semi-log plot and obtain the slope of the line, as the population growth rate. In order to improve the estimate and make use of the entire dataset, we take 100 permutations of the order of the cells during population growth and compute the growth rate. We take the average from the 100 estimates as the population growth rate.

### 6 Fitting to Log-GEV distribution

In a log-GEV distribution log transformed variables follow a GEV distribution. Thus, we plot the histogram of the log transformed generation times, and then fit a GEV distribution to the log transformed dataset to obtain the fitted Probability Density Function (PDF). We note the edges of the histogram bins in log scale and transform them back to the linear scale to obtain the log-GEV distribution.

### 7 Statistical Tests

We have performed 2-way ANOVA tests using *anova2* in MATLAB on the datasets comparing the effect of multiple parameters on the growth rate in the manuscript, to test whether the observed patterns are statistically significant.

## Acknowledgements

We are grateful to Christopher Marx for valuable discussions and feedback.

The authors also acknowledge the ICTS-ICTP Quantitative Systems Biology Schools 2015 and 2017 and the ICTS Bangalore School on Population Genetics and Evolution 2020, for the learning opportunity and exposure they provided.

D.D. was supported by funding from the Department of Science & Technology, Innovation in Science Pursuit for Inspired Research (DST-INSPIRE) Fellowship (IF160091), Government of India.

We declare that we have no competing interests.

D.D. and S.S. conceptualised the study. D.D. and S.S. developed the theory. D.D. performed the simulations. D.D. wrote the manuscript with support from S.S.

