## Supplementary Information for "Cell growth model with stochastic gene expression helps understand the growth advantage of metabolic exchange and auxotrophy"

### S1. Analysis of Distributions of generation times obtained from simulation

Here we plot the Coefficient of Variation (CV), Skewness, Kurtosis, for the distributions of generation times obtained in various simulated conditions.

We also plot the log GEV fit parameters – Shape (k), Location (μ), Scale (σ).

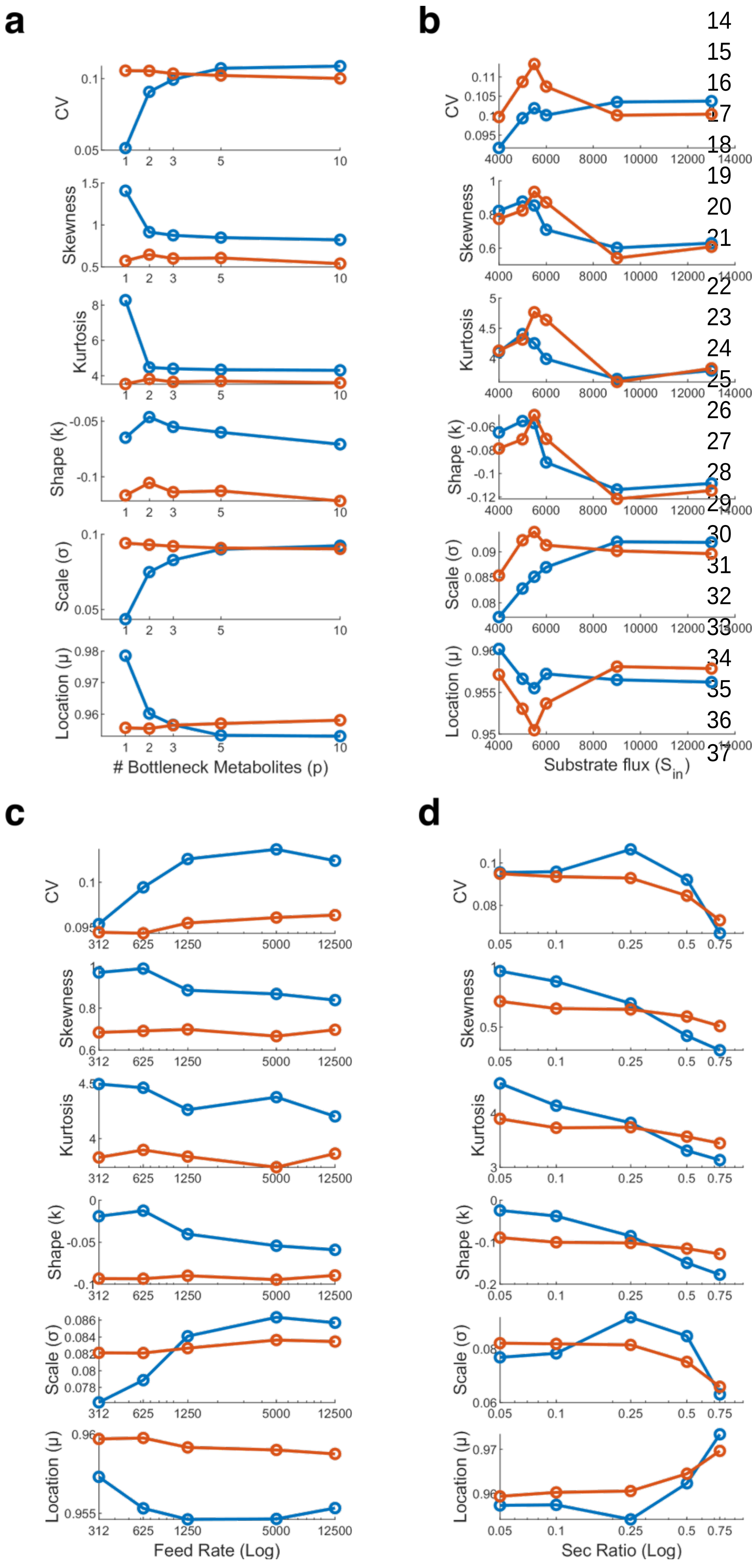

Figure S1: Variation of CV, Skewness, Kurtosis, and the log GEV Fit parameters for distributions of generation times across different conditions

(a) Varying p (Fig 2b), Low Sin (blue), high Sin (red).

(b) Varying Sin (Fig 2c), blue – p=3, red – p = 10.

(c) Varying metabolite import/feed rate (Fig 3a and 3b), low Sin (blue), high Sin (red).

(d) Varying metabolite secretion ratio (Fig 3e and 3f), low Sin (blue), high Sin (red).

S2. Analysis of experimentally observed single cell growth of *E. coli* in different media conditions (Dataset from Taher-Araghi *et al*. Curr Biol 2015)

Growth of E. coli at a single cell resolution at various media conditions was studied using microfluidic device called Mother machine, in (1).

They measured the growth of E. coli in the following media conditions: Tryptone Soya Broth (TSB), Synthetic Rich media, glucose, glucose+6aa, glucose+12aa, sorbitol, and glycerol.

We analysed the dataset to obtain the cell doubling times and plotted the distributions. We fit the distributions to both Generalized Extreme Value distributions (Fig – S2a) and Log-Generalized Extreme Value distributions (Fig – S2b).

In order to compare the fit of the data between these two distributions, we sampled 1e6 random data points from the fit distributions of the two types, and then ran a two sampled Kolmogorov-Smirnov test, with the original dataset and the sampled random data from the two distributions, and found the test was unable to reject the null hypothesis that both datasets are from the same distribution, even at 1e–8% significance level! Thus, from our analysis we find both types of distributions (GEV and log-GEV) with appropriate parameters can fit the bacterial cell generation time distributions well.

We plot the fit parameters of the GEV distribution and log-GEV distribution across different media conditions from the dataset in Figure – S2c and S2d respectively. We find that the fit value of the shape parameter (k) < 0, across all the conditions in the dataset. Thus, our analysis finds the distributions fit best by the Reverse Weibull (GEV Type III) and log-Reverse Weibull, and not by log- Fréchet (GEV Type II, k > 0) as found by previous analysis (2). The difference in our results could stem from the fact that the dataset we analyse is different. Additionally, the magnitude of fit value of shape parameter is very low (GEV: -0.07 to -0.02 and log GEV: -0.15 to -0.05), thus the nature of the shape is not too different from Fréchet with shape parameter k > 0.

We also analysed the dataset to obtain its mean generation time, the coefficient of variance (CV), skewness and kurtosis (Fig – S2e). The dataset shows that the CV of the experimentally obtained distribution ranges from about 6–14%.

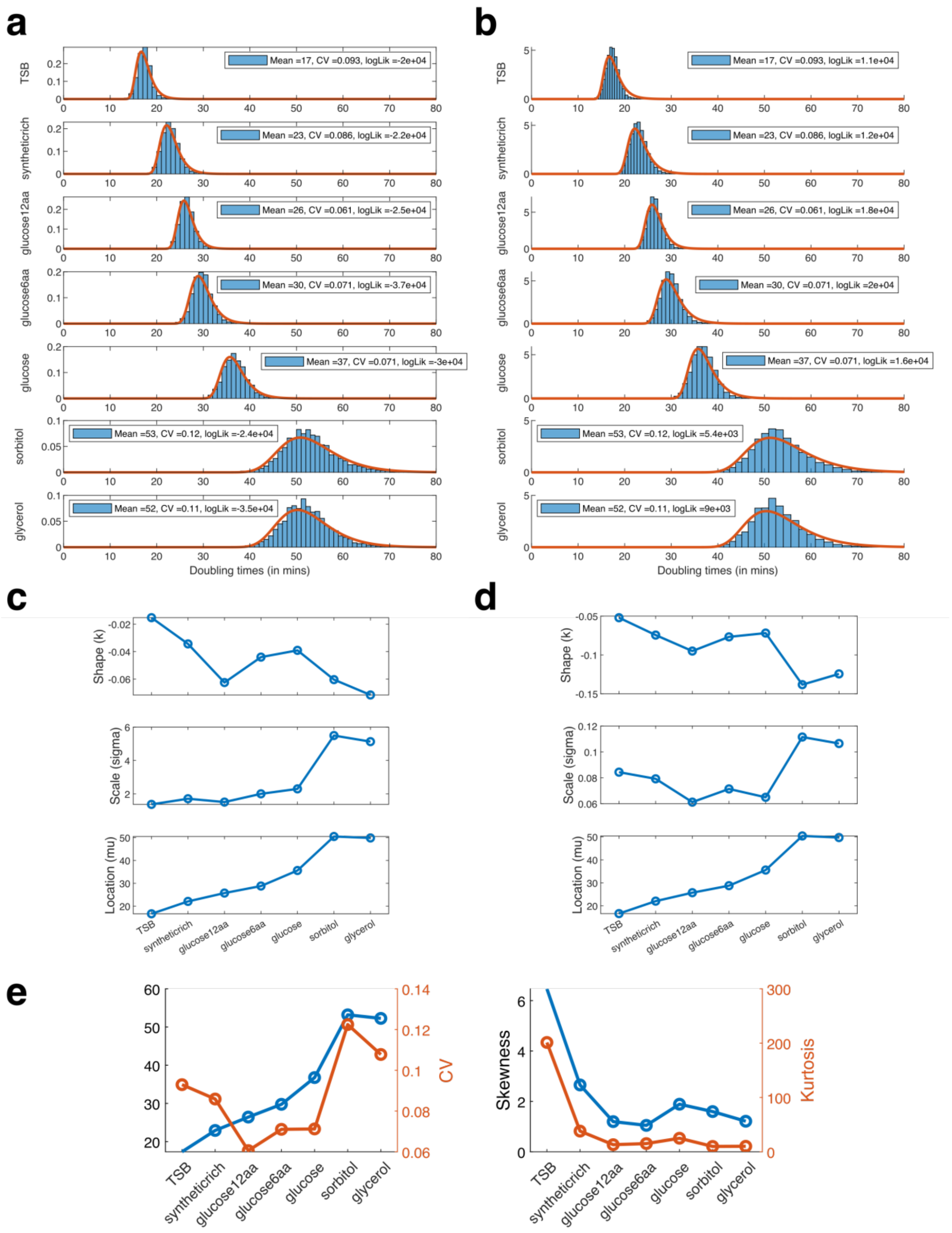

Figure S2: Analysis of the Taher-Araghi 2015 dataset

(a) Fit of the dataset to GEV distribution. (b) Fit of the dataset to log-GEV distribution. Fitted parameters of (c) GEV distribution, (d) log-GEV distribution, for each of the different media conditions in the dataset arranged in ascending order of mean generation time.

### S3. Shape of the distribution of bacterial doubling times from our model

We started our model using the original parameters of $t_{ON}$ and $t_{OFF}$ from Golding et al. 2005, which led us to generation time distributions that had too high a Coefficient of Variation (CV ~ 0.45), when we chose a metabolite threshold value that yielded physiological mean generation times ~30-50 mins. (Fig – S3a).

Upon varying the metabolite threshold, we found that we could obtain the requisite range for the CV, but using a much higher value for the threshold (Fig – S3b), which also results in a much greater mean generation time (Fig – S3b). Moreover, the shape of the distribution also changes greatly due to the variation of the metabolite threshold parameter (Fig – S3c). However, the change in shape is reminiscent of how the shape of the GEV distribution changes as its parameters change.

Within the framework of our simplistic model, the above observation suggests that the noise profile of growth-determining biosynthetic metabolic enzymes in a bacterial cell is much lower than the noise profile of a synthetic promoter whose expression characteristics was measured in Golding et al. 2005 (3). We, hence searched the parameter space of the $t_{ON}$, $t_{OFF}$, and threshold parameters to find the values at which we can find physiological values of mean generation time as well as the CV (Fig – S3d–S3g). We finally chose: $t_{OFF}$ = 2.4 min, $t_{ON}$ = 4 min, Threshold = 1e7.

### Additional factors that affect cell generation time distribution

#### 1. DNA Replication

Our model captures the dynamics of metabolite accumulation due to stochastic gene expression, and its effect on growth. As cell grows, more proteins are made but cell size also increases, thus protein concentrations that determine the probability of burst frequencies remain unaltered. However, DNA replication also takes place during growth, which could potentially double the rate of gene expression, if not limited by the transcription machinery. In case replication increases gene expression, protein number and metabolite production also should increase, which in turn may allow the slow pathways to produce more and reach the requirement threshold earlier and hence decrease the generation times. Thus, with DNA replication incorporated, we expect the generation time distributions may reduce its width (CV) for the same set of gene expression parameter used in the model.

2. Cell size variations

Our model assumes strict metabolite level sizer and adder laws for division, which can be thought of as a proxy for cell size. In our model we have considered both perfect partition between the cells as well as imperfect partitioning (Supplementary S6), and find little difference between the nature of the outcomes. Previous models have also claimed that it is not possible to distinguish between the nature of noises from gene expression and imperfect partitioning (4).

Further in case, there is a noise in terms of the cell's size perception to trigger division, it is likely to only add an extra short delay on to the result. Additionally, experiments have shown that cells seems to follow the size law when growing in poorer media at slower rates, and follow the adder when growing faster in richer media (5). We do not understand how such changes alter the generation time distribution.

Figure – S3: Shape of the distribution of bacterial doubling times from our model

(a) Generation time distribution with Golding parameters. Change in (b) CV and mean generation time, (c) Shape of distribution, with metabolite threshold. Change in CV (d,f) and mean generation time (e,g), with changing metabolite threshold, and varying gene expression parameters: tON (d,e), tOFF (f,g).

### S4. Metabolic Adder and Metabolic Sizer Models

Our framework connects stochastic gene expression with growth by transforming the cell size based empirical laws: Adder and Sizer. We assume that any cell size change is a result of equivalent quantity of metabolites produced in proper stoichiometry, and propose that cell divides upon having produced the requisite quantities of the metabolites.

In case of metabolic Adder, whatever be the existing number of metabolites in the cell after birth, the cell needs to produce the new metabolites equal to the threshold parameter. Whereas, in case the Sizer, the cell needs to only produce the difference in quantity between the existing and the threshold (Fig – 1a). We consider that the cell divides only when all the metabolites cross their respective set thresholds, in keeping with a strict requirement of stoichiometry.

We compare the outcome of the generation time distribution when we implement metabolic Adder vs the metabolic Sizer in our model, in Fig – S4a. We found that the distributions obtained for adder and sizer are pretty similar, although they tend to diverge as the number of concurrent cascades ($p$ increases), and also when the substrate import flux ($S_{in}$) is low.

When we compare the distributions of generation times from sizer and adder after rescaling the distributions with the mean, we can see that adder distributions are narrower with a lower CV (Fig – S4b).

For a single metabolic bottleneck, both laws give identical results (Fig – S4a). However, increase in the number of bottlenecks leads to a deviation between the two models, where, the Adder law consistently yields higher mean generation times. We chose to go ahead with Sizer for all the subsequent simulations, since it is reported that for poor media conditions, i.e. growth on single substrates in minimal media, the observed results suit a sizer better. While the results of growth in rich media, suit an adder better (5).

Lastly, both adder and sizer based simulations result in distributions of generation times that are right skewed and long tailed and fit well by Log GEV distributions (Fig – S4c and S4c).

Figure S4: Comparison of Metabolic Adder and Metabolic Sizer Models. Distributions of generation times from adder and sizer compared, with the mean value (a), rescaled distributions with the CV (b). Fit of the generation time distribution to log-GEV distribution (c) Sizer, (d) Adder.

### S5. A simplified steady-state version of reciprocal cross-feeding model

Our model implements the idea of a “metabolic adder” or “metabolic sizer” for cell division, along with a stochastic framework, to reveal the effect of noise on growth outcomes. However, the same model considerations when implemented as a steady state model, cannot distinguish any differences in growth between the different cases.

Take for example, the case of prototroph, with 3 concurrent limiting metabolites ($p = 3$), each of which is produced by a 3-step linear enzyme pathway ($n = 3$). In accordance with a steady-state understanding, we consider a fixed enzyme amount present in the cell. The model is fed by the constant inflow of the substrate and cell divides when the threshold requirement of each of the metabolites is met. The time duration between successive divisions is noted. Post division the cellular contents are halved and the process repeats.

We find that while we use this steady state model, at higher substrate flux values the prototroph and the auxotroph’s mean generation times are the same (Fig – S5). However, at low substrate flux rates, the auxotrophs are able to yield a shorter generation time, based on the rate of import of the missing metabolite. The advantage at low flux rates come from the fact that deletion of the pathway, allow for the substrate flux originally intended for 3 pathways are not used only by 2, which combined with a fast rate of missing metabolite import, can reduces the time required for division.

However, in the case, when the same auxotroph now directly uses the saved enzyme producing resource from one pathway to double the amount of enzyme for another pathway, then we see that the prototroph and the secreting auxotroph has no difference in their mean generation times.

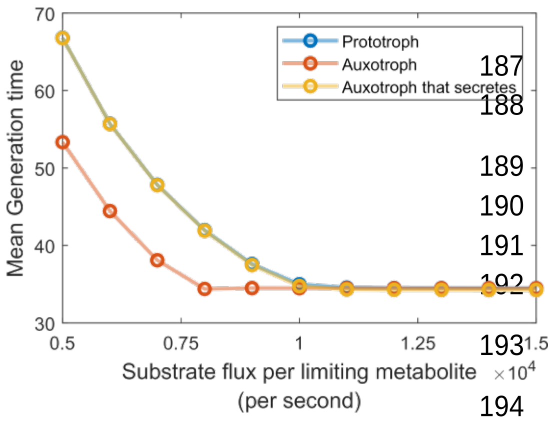

Figure S5: A simplified steady-state version of reciprocal cross-feeding model

Comparing the mean generation time of cells, when considering a steady-state model of prototrophs, auxotrophs and auxotrophs that overexpress and secrete.

### S6. Effect of different modelling criteria on the nature of cell generation times observed

#### Imperfect partition of cells

The only source of stochastic noise we consider in our model is the burst-like gene expression process. However, other sources can also add noise to the cellular growth outcome such as variation in cellular partition ratio, protein translation, and mRNA degradation. Nevertheless, their relative importance varies across different growth rates, with gene transcription and cellular partition being the most important, while mRNA degradation and protein translation are important only at very low rates of growth (6). Hence, we discuss only the effect of imprecise partitioning.

Although the process of cell division in bacteria is highly robust and precise, the two daughter cells obtained are not always of equal sizes. Discrepancies in partition ratio are rare in simple rod-shaped cells, but higher in case of other cell shapes, and on an average, the Coefficient of Variation (CV) in terms of cell length ranges from 3 – 15% (7). Moreover, the profile of noise contribution due to unequal partitioning of the cell has been shown to be virtually indistinguishable compared to stochastic gene transcription (4). We incorporate a randomly sampled partition ratio from a Normal distribution, with mean 0.5 and CV = 0.1, to simulate imperfect division. We observe that the additional noise delays the cell division process, and the distribution of generation times is shifted to the right (Fig – S6).

#### Effect of gene organisation as Operon

Bacterial genes that are co-expressed are mostly organised as a contiguous unit in the genome called operon (8–13). Our model design thus assumes that all the biosynthetic pathway enzymes occur in the same operon and share the same transcriptional burst profile. If the enzymes were organised separately and transcribed independently, each with its own burst profile, then the noisy lack of synchrony would not only lead to wasted resources (14, 15), we also expect a delay in the metabolite production and hence cell division. Simulating our model with independent expression shows the distribution of generation times is shifted to the right, towards higher values (Fig – S6).

Our model thus demonstrates that operon structure may also lead to kinetic (growth rate) gains for the cell, in addition to the already hypothesised economic gains (14).

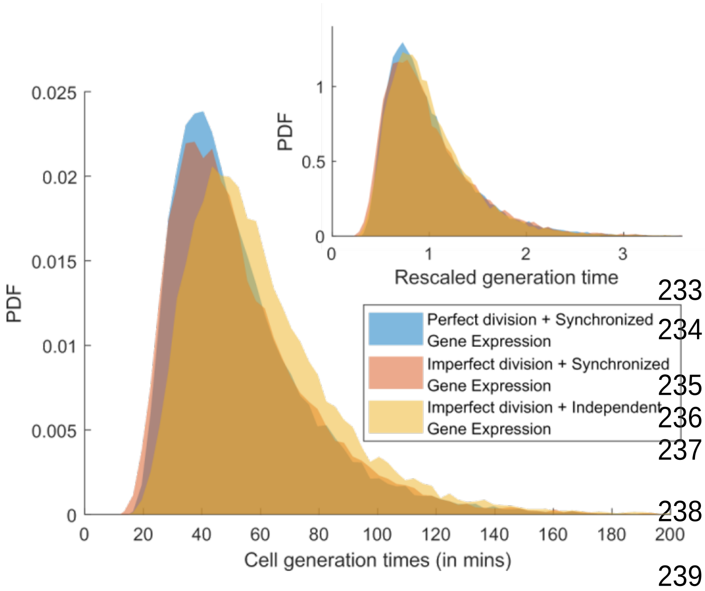

Figure S6: Effect of different modelling criteria on the cell generation time distributions

(Inset) shows the distributions rescaled by mean.

<u>Note</u>: The results shown here use a different set of gene expression and metabolite threshold than in the main paper.

### S7. Statistical Test to validate the observed patterns in the data

Here we report the Statistical tests done to show the statistical significance of the patterns we observe.

We used Two-way ANOVA to compare the effect of two parameters simultaneously on the growth rate.

We used anova2 in MATLAB, to perform the test and across conditions, our simulation results report that the tested groups are unequal from each other with a probability reported as $p = 0$ (i.e. $p < 2E\text{-}272$). However, the test also reports very high interaction between the two parameter groups compared, which is expected.

<u>Table headers</u>: Sources of variability (Source), Sum of squares due to each source (SS), Degrees of freedom associated with each source (df), Mean squares for each source, which is the ratio SS/df (MS), F-statistic, which is the ratio of the mean squares (F), p-value, which is the probability that the F-statistic can take a value larger than the computed test-statistic value (Prob>F).

#### Table for Fig – 2d

| Source | SS | df | MS | F | Prob>F |
|---|---|---|---|---|---|
| Columns | 4.65E+06 | 4.00E+00 | 1.16E+06 | 7.89E+04 | 0.00E+00 |
| Rows | 8.70E+08 | 1.40E+01 | 6.22E+07 | 4.22E+06 | 0.00E+00 |
| Interaction | 1.67E+06 | 5.60E+01 | 2.98E+04 | 2.03E+03 | 0.00E+00 |
| Error | 2.71E+07 | 1.84E+06 | 1.47E+01 | | |
| Total | 9.04E+08 | 1.84E+06 | | | |

#### Table for Fig – 3c

| Source | SS | df | MS | F | Prob>F |
|---|---|---|---|---|---|
| Columns | 3.64E+08 | 4.00E+00 | 9.10E+07 | 2.56E+03 | 0.00E+00 |
| Rows | 1.77E+10 | 5.00E+00 | 3.55E+09 | 9.98E+04 | 0.00E+00 |
| Interaction | 4.78E+07 | 2.00E+01 | 2.39E+06 | 6.72E+01 | 1.53E-272 |
| Error | 6.11E+10 | 1.72E+06 | 3.55E+04 | | |
| Total | 7.93E+10 | 1.72E+06 | | | |

Table for Fig – 3g

| Source | SS | df | MS | F | Prob>F |
|---|---|---|---|---|---|
| Columns | 1.62E+12 | 5.00E+00 | 3.23E+11 | 4.41E+06 | 0.00E+00 |
| Rows | 1.48E+11 | 5.00E+00 | 2.95E+10 | 4.02E+05 | 0.00E+00 |
| Interaction | 2.17E+11 | 2.50E+01 | 8.69E+09 | 1.18E+05 | 0.00E+00 |
| Error | 6.49E+10 | 8.85E+05 | 7.34E+04 | | |
| Total | 2.05E+12 | 8.85E+05 | | | |

Table for Fig – 4b

Compared only the results of simulating auxotroph growth (solid colour lines in Fig – 4b), and neglected feed = 12500, since the growth rate saturates and feed = 7500 and 12500 are identical.

| Source | SS | df | MS | F | Prob>F |
|---|---|---|---|---|---|
| Columns | 1.88E+05 | 5.00E+00 | 3.76E+04 | 2.97E+00 | 0.011 |
| Rows | 5.47E+11 | 2.00E+00 | 2.74E+11 | 2.16E+07 | 0.000 |
| Interaction | 2.55E+05 | 1.00E+01 | 2.55E+04 | 2.01E+00 | 0.028 |
| Error | 7.47E+09 | 5.90E+05 | 1.27E+04 | | |
| Total | 5.55E+11 | 5.90E+05 | | | |

Table for Fig – 5b

| Source | SS | df | MS | F | Prob>F |
|---|---|---|---|---|---|
| Columns | 1.13E+12 | 2.80E+01 | 4.03E+10 | 1.39E+06 | 0.00E+00 |
| Rows | 9.81E+10 | 4.00E+00 | 2.45E+10 | 8.45E+05 | 0.00E+00 |
| Interaction | 4.72E+10 | 1.12E+02 | 4.22E+08 | 1.45E+04 | 0.00E+00 |
| Error | 1.38E+11 | 4.75E+06 | 2.90E+04 | | |
| Total | 1.41E+12 | 4.75E+06 | | | |

Table for Fig – 6a

| Source | SS | df | MS | F | Prob>F |
|---|---|---|---|---|---|
| Columns | 1.74E+08 | 6.00E+00 | 2.89E+07 | 8.11E+02 | 0.00E+00 |
| Rows | 3.60E+09 | 5.00E+00 | 7.19E+08 | 2.02E+04 | 0.00E+00 |
| Interaction | 5.91E+08 | 3.00E+01 | 1.97E+07 | 5.53E+02 | 0.00E+00 |
| Error | 2.45E+10 | 6.88E+05 | 3.56E+04 | | |
| Total | 2.89E+10 | 6.88E+05 | | | |

Table for Fig – 6b

| Source | SS | df | MS | F | Prob>F |
|---|---|---|---|---|---|
| Columns | 2.00E+10 | 7.00E+00 | 2.86E+09 | 7.76E+04 | 0.00E+00 |
| Rows | 4.12E+10 | 4.00E+00 | 1.03E+10 | 2.80E+05 | 0.00E+00 |
| Interaction | 3.75E+09 | 2.80E+01 | 1.34E+08 | 3.64E+03 | 0.00E+00 |

| | | | |
|---|---|---|---|
| Error | 4.82E+10 | 1.31E+06 | 3.68E+04 |
| Total | 1.13E+11 | 1.31E+06 | |

Table for Fig – 6c

| Source | SS | df | MS | F | Prob>F |
|---|---|---|---|---|---|
| Columns | 4.53E+09 | 4.00E+00 | 1.13E+09 | 2.99E+04 | 0.00E+00 |
| Rows | 2.68E+10 | 4.00E+00 | 6.69E+09 | 1.77E+05 | 0.00E+00 |
| Interaction | 1.09E+09 | 1.60E+01 | 6.83E+07 | 1.80E+03 | 0.00E+00 |
| Error | 3.10E+10 | 8.19E+05 | 3.79E+04 | | |
| Total | 6.34E+10 | 8.19E+05 | | | |

Table for Fig – 6d

| Source | SS | df | MS | F | Prob>F |
|---|---|---|---|---|---|
| Columns | 5.01E+09 | 5.00E+00 | 1.00E+09 | 3.58E+04 | 0.00E+00 |
| Rows | 4.66E+09 | 4.00E+00 | 1.16E+09 | 4.16E+04 | 0.00E+00 |
| Interaction | 2.20E+08 | 2.00E+01 | 1.10E+07 | 3.93E+02 | 0.00E+00 |
| Error | 2.75E+10 | 9.83E+05 | 2.80E+04 | | |
| Total | 3.74E+10 | 9.83E+05 | | | |

### S8. Comparison of generation time distributions of prototrophs and auxotrophs when growth rates are equal

Here we compare the overlap of the distributions of generation times between cases when the growth rates computed from the generation times are equal. In our simulations we demonstrate that by externally supplying a limiting metabolite, prototrophs can grow faster, same as the prototroph with one less bottleneck. Auxotrophs can also enjoy the same benefit by losing one of the limiting metabolite biosynthetic pathway.

We compare between the generation time distribution of a prototroph with two bottleneck metabolites ($p = 2$), against the following: (a) prototroph ($p = 3$), fed one limiting metabolite at a high rate, (b) auxotroph (originally $p = 3$), fed the missing limiting metabolite at a high rate.

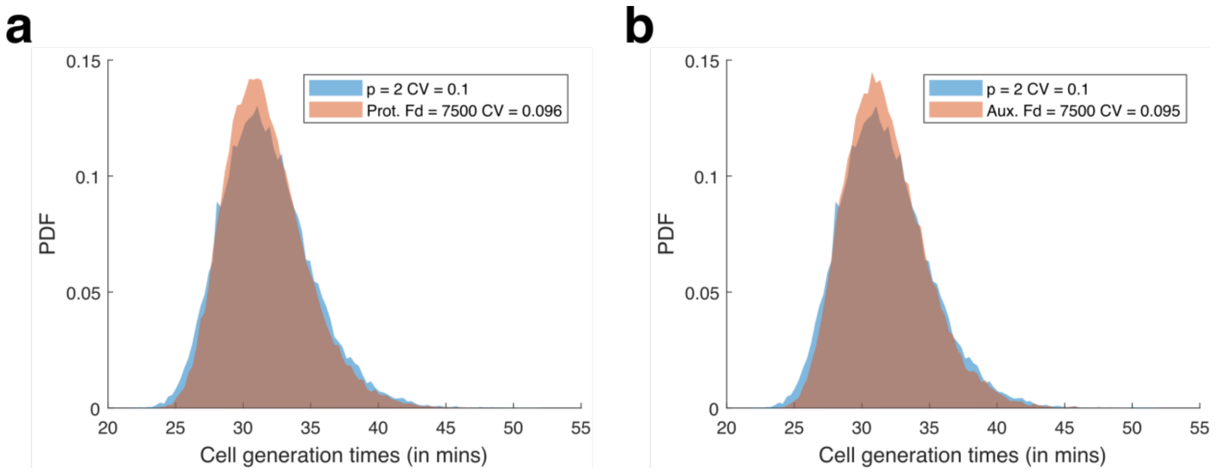

Figure S8: Comparison of generation time distributions of prototroph with lower bottleneck (p = 2) versus (a) prototroph, and (b) auxotroph, when fed the limiting metabolite at 7500/sec such that the growth rates are equal.

S9. List of Parameters

| PARAMETER | VALUE | SOURCE |
|---|---|---|
| Mean mRNA lifetime | 25 mins | Bionumbers 112326 |
| Mean protein lifetime | 20 hours | Bionumbers 111930 |
| Average Enzyme gene length | 1029 bp / 343 aa | Bionumbers 108985 |
| Transcription Rate | 55 nt/sec | Bionumbers 100059 |
| Translation Rate | 20 aa/sec | Bionumbers 100059 |
| Mean duration of a Transcription burst ($\boldsymbol{t_{ON}}$) | 4 mins | Optimized (See Supplementary S3) |
| Mean wait time duration between Transcription bursts ($\boldsymbol{t_{OFF}}$) | 2.4 mins | Optimized (See Supplementary S3) |
| Median enzyme turnover number or catalytic constant ($\boldsymbol{k_{cat}}$) | 10 per sec | Bionumbers 111411 |
| Median enzyme half-saturation constant ($\boldsymbol{K_M}$) | 100 μM or 60220 molecules | Bionumbers 111413 (Value converted to molecules assuming a cell volume of 1μm$^3$) |
